# Root production in a subtropical pasture is mediated by cultivar and defoliation severity

**DOI:** 10.1101/763128

**Authors:** Chris H. Wilson, Joao M. Vendramini, Lynn E. Sollenberger, S. Luke Flory

## Abstract

**Background:** Grasslands occupy significant land area and account for a large proportion of the global soil carbon stock, yet the direct effects of grazing and genotypic composition on relationships between shoot and root production are poorly resolved. This lack of understanding hinders the development of models for predicting root production in managed grasslands, a critical variable for determining soil carbon stocks.

**Methods:** We quantified the effects of season-long defoliation treatments on both shoot and root production across four cultivars of a widely-planted pasture grass species (*Paspalum notatum* Fluegge) in a common garden setting in South Florida, USA.

**Results:** We found that infrequently applied (4 week) severe defoliation (to 5 cm) substantially enhanced shoot production for all cultivars, while severe defoliation reduced root production across cultivars, regardless of frequency. Overall, cultivars varied substantially in root production across the range of defoliation treatments in our study. However, there was no significant relationship between shoot and root production.

**Conclusions:** Our results find that aboveground and belowground productivity are only weakly coupled, suggesting caution against use of simple aboveground proxies to predict variations in root production in grasslands. More broadly, our results demonstrate that improved modeling and management of grasslands for belowground ecosystem services, including soil carbon sequestration/stocks, will need to account for intraspecific genetic variations and responses to defoliation management.

## Introduction

Grassland ecosystems occupy more than a fifth of earth’s land area and account for a large proportion of the global SOC stock (1,2). However, there is considerable uncertainty in predictions of net ecosystem exchange, and hence carbon sequestration services from grasslands (3,4). One significant source of uncertainty is that while large herbivore grazing is known to mediate patterns of plant species composition, diversity, and aboveground primary productivity (5–7), the effects of grazing on belowground processes and soil carbon is less clear (8–11). In particular, there are limited field studies where the impact of grazing on root production in grassland systems has been directly measured (e.g., via root ingrowth cores or minirhizotron technology, but see Ziter and MacDougall (12) Balogianni et al. (11) and Cooley et al. (13)). Since belowground production may be the largest component of total NPP for many grasslands (14,15), determining how grazing affects root production will help to predict if and when grassland ecosystems will behave as carbon sinks, and whether grazing is likely to promote or inhibit carbon sequestration services.

Root carbon inputs may constitute a disproportionate amount of the total SOC stock compared with shoot carbon (16–18), and are especially critical in grassland ecosystems where aboveground tissue is susceptible to frequent removal by fire and grazing (19). Current understanding of how grazing affects root production is ambiguous. For example, one temperate mesocosm study showed that intense defoliation inhibited root production and accelerated the loss of SOC (20), whereas some field studies have documented greater belowground allocation and root production under grazing (Hafner et al. (21) in the Tibetan plateau; Wilson et al. (22) in subtropical pasture).. Augustine et al. (23) found that defoliation reduced belowground carbon allocation in one grazing-adapted North American grass species (*Pascopyrum smithii*, western wheatgrass) but not in another (*Bouteloa gracilis*, blue grama), highlighting interspecific variations in response to a given defoliation regime. In general, laboratory and mesocosm studies have found that frequent grazing/defoliation leads to declines in standing root biomass over the long term (24), whereas a global synthesis of data comparing grazed and ungrazed grasslands found a mix of positive and negative effects on standing root biomass (25). Overall, this discordance suggests that variations in plant composition, underlying environmental factors, grazing intensity, or some combination of these factors significantly mediate the effect of grazing on root production.

Grazing effects on belowground production may not only vary based on plant species, but also on the genotypic composition of a grazed stand, given the increasing evidence of the importance of intraspecific variation in driving ecosystem structure and function (26,27). In general, some literature suggests that reduced root allocation (and increased shoot allocation) following grazing may represent an evolutionarily adaptive trait for grazing tolerance (28). For instance, Carman (1985) (29) noted that short-leaved genotypes of *Schizachyrium scoparium*, selected from a long-term grazed site, exhibited lower rates of root elongation post-grazing than longer-leaved genotypes from a long-term grazing excluded site. Planted pasture grasses also have been shown to exhibit genotypic variability in shoot and root production in response to grazing (e.g. Dawson et al. (30)). For example, Interrante et al. (31) observed significantly less plant cover in recently selected, upright-growing *Paspalum notatum* (bahiagrass) cultivars in response to severe, frequent defoliation, but did not observe less cover with the same defoliation treatments applied to widely naturalized cultivars, suggesting significant intraspecific variability in grazing tolerance and belowground allocation.

Although root production is a critical component of predicting the carbon cycle in grassland ecosystems, it is difficult to monitor or predict over large spatial scales. Thus, regional-scale grassland models have been developed that predict total NPP and/or greenhouse gas exchange on the basis of aboveground canopy characteristics estimated from remote sensing (32–34). Similarly, some previous work has sought to predict BNPP on the basis of readily obtained aboveground measurements in both grasslands (14) and forests (35). Recently, concerted efforts have been made to link fine root traits with other plant traits, across species and environments, by compiling and analyzing global-scale big datasets (36). The goal is to have reliable aboveground proxies for predicting critical belowground root processes (37). However, given the evidence for potentially significant genotypic and defoliation effects on belowground carbon allocation, it is unclear whether aboveground proxies can ever reliably approximate root production. Given the central importance of root system carbon inputs to maintaining SOC, especially in grasslands, we need more data from experimental systems where genotypic composition and grazing management have been manipulated, and the relationship between above and belowground allocation have been quantified.

In this study, we tested the independent and combined effects of defoliation intensity and frequency, and cultivar on root production of a widely utilized pasture grass species of the southeastern United States, *Paspalum notatum* Flüegge (bahiagrass). For Bahiagrass, we can broadly delineate cultivars on the basis of growth habit where historically older, widely naturalized cultivars tend to have a more prostrate growth pattern, whereas recently selected cultivars tend to have a more upright growth pattern, reflecting selection for improved forage growth characteristics (38). Previous work, and considerable producer experience, suggests that bahiagrass has a remarkable resilience to intense grazing, wherein forage growth and quality is maximized with severe defoliation (close to ground level) so long as regrowth intervals are adequate (39,40). However, the impact of defoliation severity on root production across cultivars, and their associated growth habits, has not been directly studied, reflecting a general lack of information on belowground growth responses in warm season subtropical pasture (13). To redress this gap in knowledge we conducted an experiment in a common garden setting under realistic conditions of limited soil fertility to isolate the effects of defoliation intensity, frequency and cultivar on belowground production, and to evaluate the relationship between aboveground and belowground growth.

Consistent with the literature on compensatory growth responses from natural and planted pastures (40–42), and the literature on genotypic variability (e.g. Dawson et al. (30)) we hypothesized that:

1. Severe defoliation, applied infrequently, would stimulate increases in aboveground primary productivity (via compensatory response mechanisms), but would have neutral effects on root productivity across all cultivars;
2. Severe defoliation, applied frequently, would significantly suppress root production across all cultivars as a consequence of plant requirements to prioritize photosynthate allocation to regrowing shoots;
3. Widely naturalized, decumbent cultivars would show proportionally greater reductions in root production under severe defoliation compared to the more upright cultivars, reflecting a beneficial adaptation for increased shoot allocation following severe defoliation events; and
4. Despite alterations to belowground allocation on the basis of cultivar and defoliation treatment, shoot production and root production would positively correlate at the plot level reflecting variations in underlying soil factors determining total productivity.

## Materials & Methods

To evaluate the independent and potential interactive effects of defoliation intensity and plant cultivar on root production, we established 32 3 m x 7 m experimental plots at the University of Florida Range Cattle Research and Education Center, Ona, FL (27°26’ N, 82°55’W) in 2009. The soils were uniform and classified as Pomona fine sand (sandy, siliceous, hyperthermic Ultic Alaquod). First, we seeded plots with one of four bahiagrass cultivars (Argentine, Pensacola, Tifton-9, and UF-Riata). Bahiagrassis a perennial C4 pasture grass with improved germplasm that was introduced to Florida in the 1920s from South America and constitutes the primary forage for the Florida cow-calf industry (43). ‘Argentine’ and ‘Pensacola’ are widely-distributed, naturalized cultivars in the state of Florida with a decumbent growth habit, whereas ‘Tifton-9’ and ‘UF-Riata’ are recently-released cultivars selected for improved agronomic characteristics including more upright growth habits and less photoperiod sensitivity (31,38). Plots were fully established by the onset of the 2010 summer growing season with complete, uniform plant cover. More details, including soil fertility characteristics can be found in Vendramini et al. (38). Site weather data for this period were accessed from the Florida Automated Weather Network (FAWN, http://fawn.ifas.ufl.edu/data/), including temperature, precipitation, and evapotranspiration, and all fell within normal ranges (Table A1).

We initiated defoliation treatments on June 13^th^ 2013 and concluded field sampling 16 weeks later on October 5^th^ 2013. Although we did not measure soil moisture, the soils were all visibly waterlogged from July until the end of the experiment, as is typical in Florida Spodosol soils (43). We therefore assumed that plant growth was not limited by water availability during the sampling period, or at the very least that water availability was essentially constant across plots. Each plot (n = 32) was randomly assigned to either a frequent (2 week) or infrequent (4 week) defoliation treatment to simulate grazing stress. Each plot was divided in half and received two defoliation intensities (a severe defoliation to 5 cm residual height, and a mild defoliation to 15 cm residual height) resulting in n = 64 experimental units (Figure A1). Thus, our design was effectively split-plot with two main-plot treatments (cultivar and defoliation frequency), while our subplot factor was defoliation intensity. Overall, each cultivar X defoliation severity X defoliation frequency treatment was replicated 4 times.

**Fig. 1:**
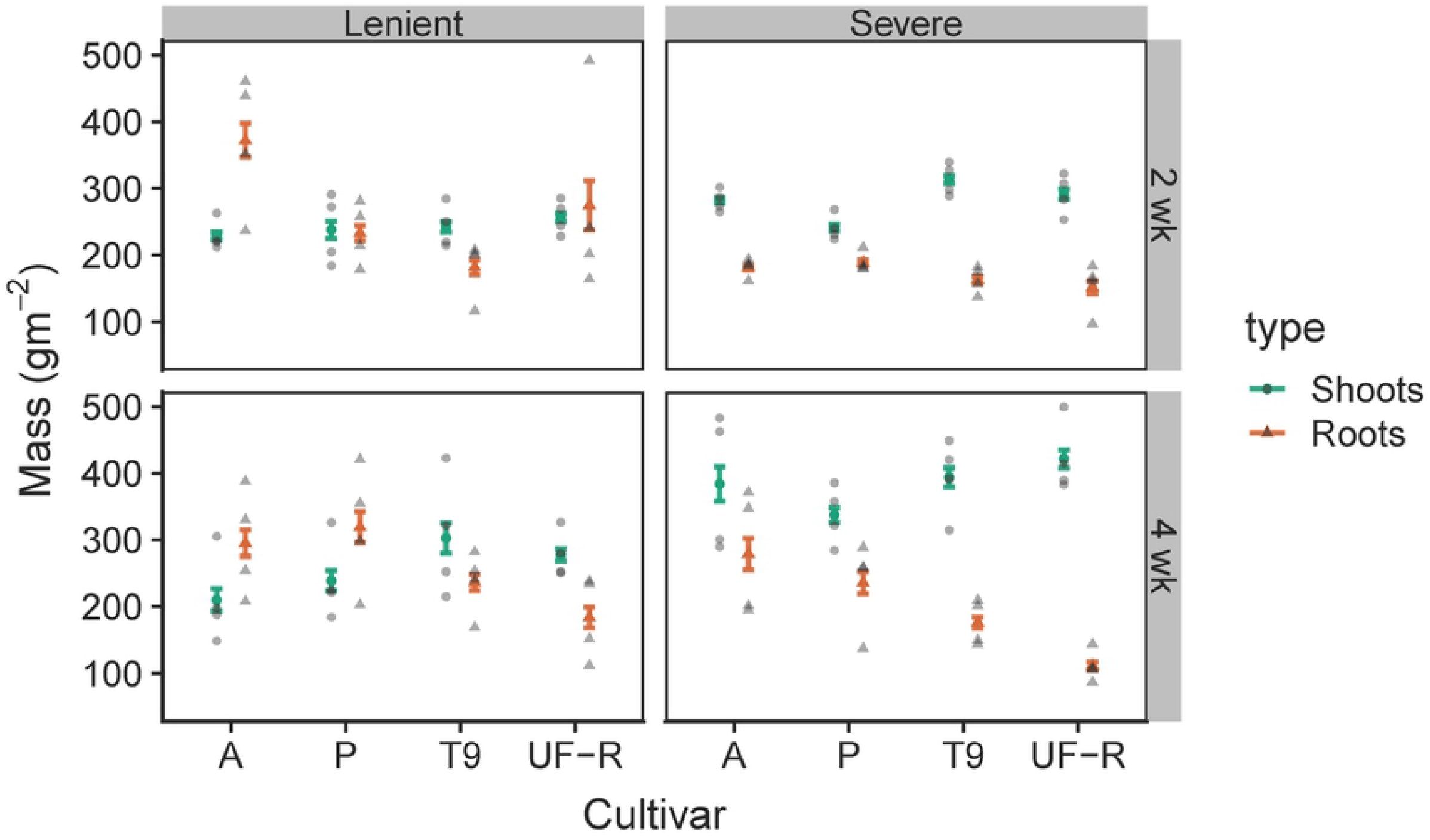
Raw data (gm^−2^) plotted as circles (shoots) and triangles (roots). Error bars show mean biomass (gm^−2^) +/- 1 SE for shoots (purple error bars) and roots (brown error bars). The panels are faceted by treatment combinations: intensity of defoliation on top (lenient 15 cm or severe 5 cm on top), and frequency of defoliation labeled on the right hand side (2 wk or 4 wk). The x-axis groups responses by cultivar: A = Argentine, P = Pensacola, T9 = Tifton-9, and UF-R = UF-Riata.

We harvested a 0.92-m^2^ quadrat from each subplot during each defoliation treatment with a rotary mower (Sensation Mow-Blo Model 11F4-0) at the target cutting heights: 5 cm for the severe defoliation, 15 cm for the mild defoliation, values chosen based on personal observation (C.H. Wilson, L.E Sollenberger, J.M. Vendramini) to represent the extremes of pasture defoliation under grazing by beef cattle in Florida. To quantify aboveground production, harvested material was oven-dried at 60°C to constant mass and weighed on an analytical scale. During the final harvest, all subplots were harvested at 5 cm. Total aboveground production was determined by summing values for each subplot across all dates including the final harvest. Aboveground production values are presented in gm^−2^ (dry biomass).

To quantify root primary production in response to the defoliation treatments, we installed 2-mm mesh root in-growth cores (44) on June 7^th^, 2013, prior to imposing the defoliation treatments. Cores were 7.5 cm diameter x 25 cm deep and constructed of fiberglass mesh. They were installed by first excavating a cylinder of soil with a soil auger to target dimensions, placing the mesh bags into the cylinder so that the upper edge of the bags was just below the soil surface, and then re-filling the cores with sieved, root-free soil from the same plot. We retrieved the cores at the end of the growing season on October 5th 2013, 16 weeks after installation. The final volume of soil contained in each core was quantified prior to washing the roots free of soil on a 250-uM sieve. Root samples were then oven-dried at 60°C to constant mass and weighed on an analytical scale. To correct for variation in core volume, root biomass was multiplied by a correction factor determined as the inverse of the ratio of each core volume to a reference core (a cylinder of 7.5 cm diameter and 25 cm depth). Finally, we visually determined that almost all root biomass was contained within the depth we evaluated (i.e. 25 cm depth) by digging several test pits around our study area. We note from personal observation that wet pastures tend to result in shallower root distribution, consistent with early literature such as (45). Therefore, we multiplied root biomass by a constant (10000/(pi*3.75^2)) to convert our measures to g/m^2^, putting them on an easily interpretable scale.

### Statistical Analysis

Response variables for analyses were shoot and root production, and a measure of root allocation defined as:

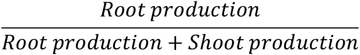

To analyze among-cultivar variability in response to our treatments, we parameterized a varying-intercept/varying-slope Bayesian hierarchical model that we applied to both of our response variables. In this model, we estimate intercept and slope (i.e., treatment effects) coefficients for each cultivar, where each batch of coefficients is modeled as a draw from a normal distribution with an estimated variance component (46). We included binary predictor variables using a −0.5/0.5 “effect coding” for our experimentally imposed treatments: lenient (15 cm) and infrequent (4 wk) defoliation were assigned −0.5 values, while frequent (2 wk) and severe (5 cm) defoliation were assigned 0.5 values. Under this coding, the model intercept represents the grand mean, and the coefficients for defoliation severity and frequency represent the main effects of severe and/or frequent defoliation across both levels of the other treatment (see e.g. Schabenberger et al. (47)). We also included a term for the interaction of severe and frequent defoliation treatments and a random effect of plot to allow for correlation in observations from the same plot. Our varying-intercept/varying-slope model therefore included four separate estimates of grand means (one for each cultivar), each of which represents an estimate of performance for that cultivar across all defoliation treatment conditions, and four treatment effect estimates (one for each cultivar) for frequent defoliation, severe defoliation, and their interaction. Since these coefficients were drawn from distributions with estimated variance components, the separate estimates were partially pooled towards their common mean, which also was estimated from the data, a property that built in an automatic correction for multiple comparisons among cultivars and obviated the need for arbitrary post-hoc adjustments such as the Bonferonni correction (48). Finally, because growth data are naturally constrained to be positive only and because we observed a pattern of variance increasing with the mean, we used a gamma distribution to model our data, which naturally accounts for this nearly universal pattern in biomass data. We used the standard log-link in our parameterization of the gamma regression model, and thus our model coefficients represent multiplicative effects, and are reported on the log link scale (46). Values greater than zero indicate positive effects on the response variable, whereas values less than zero indicate negative effects. As in all cases where the log-link is used, exponentiation of these regression coefficients returns the multiplicative effect which can be naturally interpreted as a % effect.

We display treatment effects graphically by first plotting estimated fixed effect coefficients (i.e. frequency, severity and frequency X severity) centered on the median, and include both a 50% (thick) and a 95% (thin line) uncertainty (credible) interval. These coefficients represent the overall average effects of treatment or the interaction effect across all cultivars. In addition, we graphically present the varying intercepts portion of our model, which represents the overall average deviation of each cultivar from the grand mean across all cultivars, and is thus naturally centered at zero. Here again, we include both 50% (thick) and a 95% (thin line) credible intervals. The proportion of the credible interval above or below zero can be interpreted as the Bayesian probability of that cultivar differing in response from the average across all cultivars. In the case of root allocation, we further analyze all the pairwise contrasts among cultivars (n=6 contrasts), by taking the difference between each coefficient at each iteration of the MCMC sampler. These pairwise contrasts thus represent the differences between each pair of cultivars in their overall root allocation, averaged across all treatment conditions.

We estimated these models in a Bayesian framework via Hamiltonian Monte Carlo in the packaged “rstanarm” (v2.18.2) called from R (v3.5.3) via Rstudio (v1.1.463). Prior to analysis, shoot and root production responses were standardized by dividing by their mean, resulting in this case with response variables with scale ~O(1) to facilitate faster sampling, and to help specify weakly-regularizing Normal(0,1) priors for all treatment effects. For all models we sampled the target (posterior) distribution with four chains of 2000 iterations each. Model convergence was assessed via use of the R-hat < 1.01 criterion (46) as well as by visual inspection for chain blending and stability, and monitoring of the powerful diagnostics built into rstanarm (i.e. divergent transitions and E-BFMI, citation).

To understand the relative importance of defoliation treatment and cultivar compared with shoot production for predicting root production, we first fit a simple univariate regression model using only aboveground biomass from each subplot (n=64) as a continuous covariate. We then refit our varying-intercepts/varying-slopes model while including shoot production as a continuous covariate alongside treatment and cultivar effects. We compare a Bayesian R^2^ metric between the models (49). Because the visual and R^2^ comparisons were so clear, we had no need to evaluate additional metrics of model predictive performance.

## Results

### Shoot production model

Average shoot production across all cultivars and treatment combinations in our study was 290 gm^−2^, with the highest values observed in the infrequent severe defoliation treatment, which averaged 384 gm^−2^ (Fig 1). The fixed main effect estimate (on log-link scale, and reported as posterior median +/- posterior standard error) for severe defoliation was positive [0.28 +/- 0.07, Fig 2a], while the estimate for frequent defoliation was negative [-0.18 +/- 0.08, Fig 2a]; however, the interaction was negative as well [-0.25 +/- 0.15, Fig 2a], consistent with readily observable pattern (Fig 1) that it is the combination of severe + infrequent (4 wk) defoliation that leads to over-yielding. Overall, we did not estimate substantial variability in shoot production *among* cultivars across all treatments, although the upright cultivars (UF-Riata and Tifton-9) had slightly higher production than the decumbent cultivars Argentine and Pensacola (Fig. 3a).

**Fig. 2:**
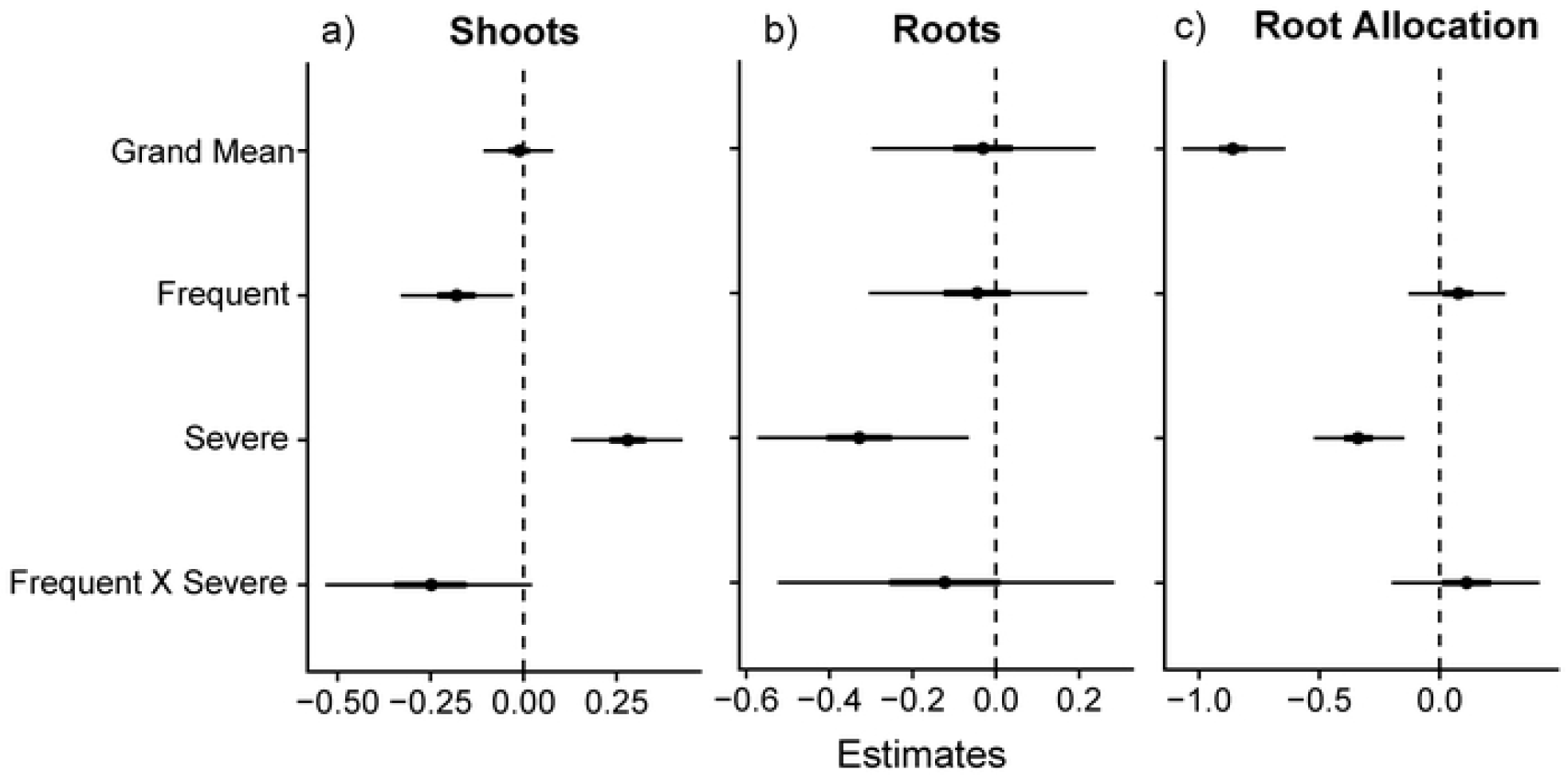
Fixed effects from varying-intercepts/varying-slopes Gamma regression model. Coefficients are plotted on the log-link scale and include a median (point), 50% (thick line) and 95% (thin line) credible intervals for a) shoot production, b) root production and c) root allocation. Where the entire 95% credible interval falls above or below zero, we can interpret that as a 97.5+% Bayesian probability of that coefficient having a positive or negative effect on the response, respectively.

**Fig. 3:**
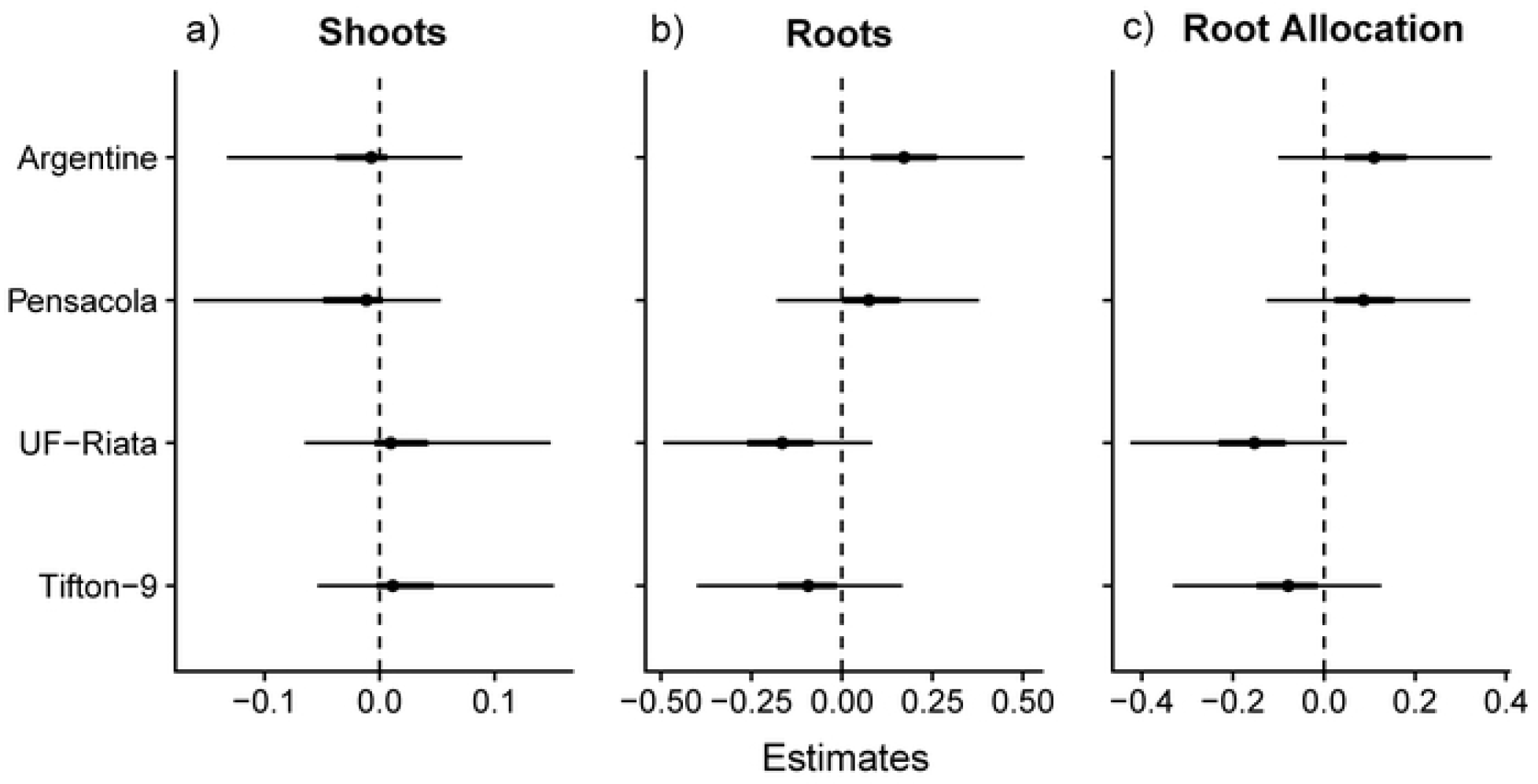
Varying-intercepts from the Gamma regression model for root production. Coefficients represent deviations of each cultivar (A = Argentine, P = Pensacola, T9 = Tifton-9, and UF-R = UF-Riata) from the overall mean (fixed effect coefficient), and are thus naturally centered at 0, where negative values represent lower than average performance, and positive values higher than average performance. Plots include a median (point), and 50% (thick line) and 95% (thin line) credible intervals. Where the entire 95% credible interval falls above or below zero, we can interpret that as a 97.5+% Bayesian probability of the cultivar having a higher or lower overall root production compared to the mean among all cultivars.

### Root production model

We observed an average root production of 224 gm^−2^, where mild defoliation treatments were the highest with 262 gm^−2^ averaged across 2 wk and 4 wk defoliation frequencies, compared with severe defoliation with an average of 186 gm^−2^ (Fig 1). The fixed main effect estimate for severe defoliation was negative (−0.33 +/- 0.12, Fig 2b), with >97.5% of posterior probability below 0, while the main effects of frequent defoliation and the interaction of frequent X severe defoliation were highly uncertain, with 95% credible intervals spanning a similar range above and below zero. Average root production across all treatment groups varied by cultivar more substantially than shoot production (Fig 3b), with the decumbent cultivars Argentine and Pensacola having greater root production than the upright cultivars UF-Riata and Tifton-9 (Fig 3b, Fig. 4). The greatest contrast was between Argentine and UF-Riata, which had a median posterior difference of −0.36 on the log-link scale (Fig. 4), which represents a 30% lower root production.

**Fig. 4:**
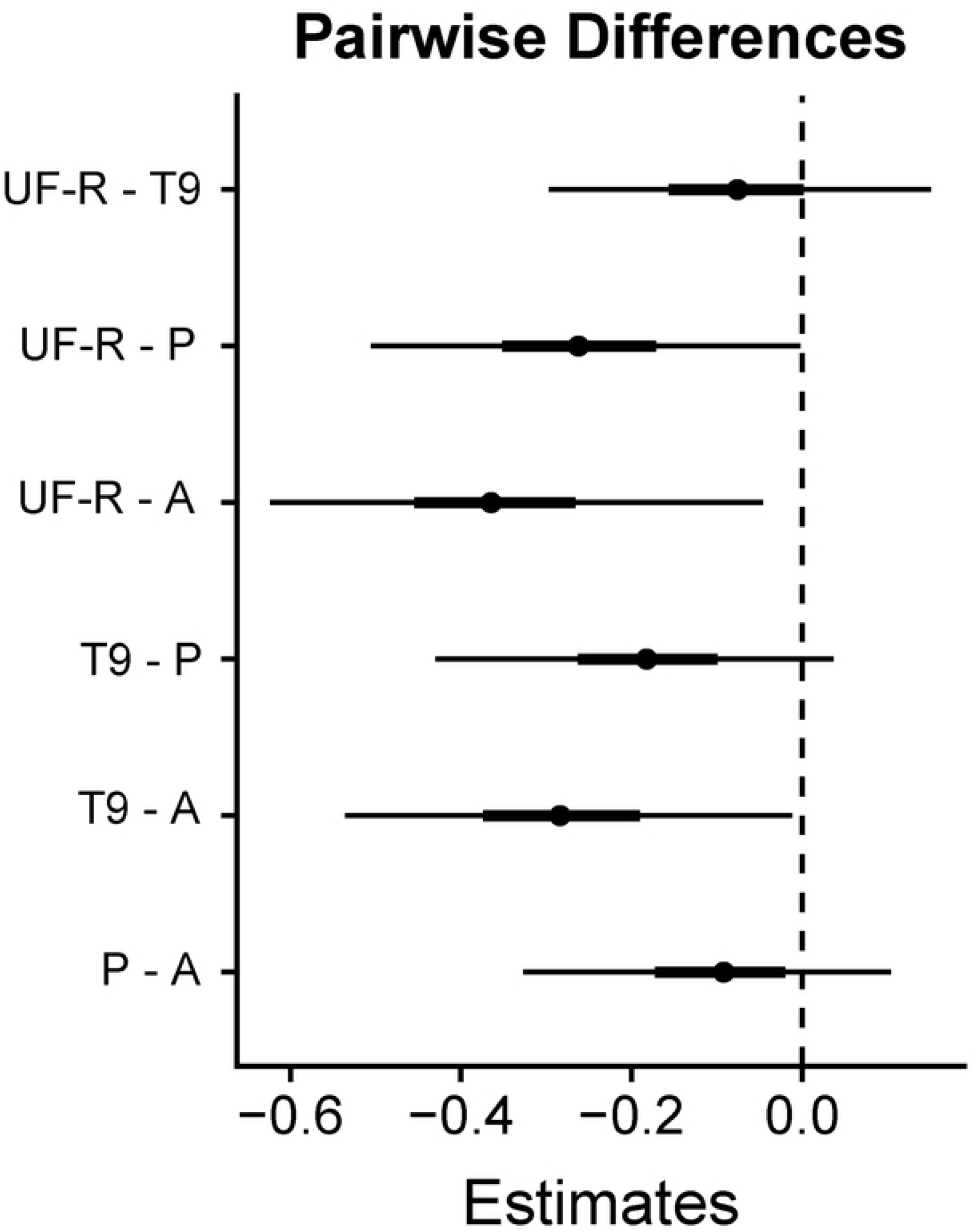
Pairwise contrasts among each cultivar for the varying intercepts of the root allocation model. Key: A = Argentine, P = Pensacola, T9 = Tifton-9, and UF-R = UF-Riata. Plots include a median (point), and 50% (thick line) and 95% (thin line) credible intervals. Where the entire 95% credible interval falls above or below zero, we can interpret that as a 97.5+% Bayesian probability of the first cultivar having a higher root allocation than the second cultivar.

### Root allocation

The fixed main effect estimate for severe defoliation on root allocation proportion was - 0.34 +/- 0.09 (Fig 2c), a very similar median estimate to that for root production, although with a smaller uncertainty (SE = 0.09 versus 0.12). This result represents a median estimate of 29% reduced allocation proportion to roots overall among cultivars and across both frequencies of defoliation with severe defoliation. Variation among cultivars was also similar to that observed for root production (Fig 3c versus 3b), and thus we did not repeat the pairwise analysis since it would convey redundant information.

### Root production predictions

The univariate regression between shoot and root production revealed a very weak (R^2^ = 0.09) relationship (Fig 5a). The full model that included treatment indicators and cultivar identity (as in the analyses above), yielded a median R^2^ of 0.45 (Fig 5b). After removing the varying intercepts/slopes by cultivar, this R^2^ value declined to 0.21 (see supplement), indicating that accounting for cultivar identity doubles model fit. Close examination of Fig 5b reveals that the full model accounted for observed variations in root production quite well in the range of 100-300 gm^−2^ but severely underpredicted root production > 300 gm^−2^.

**Fig. 5:**
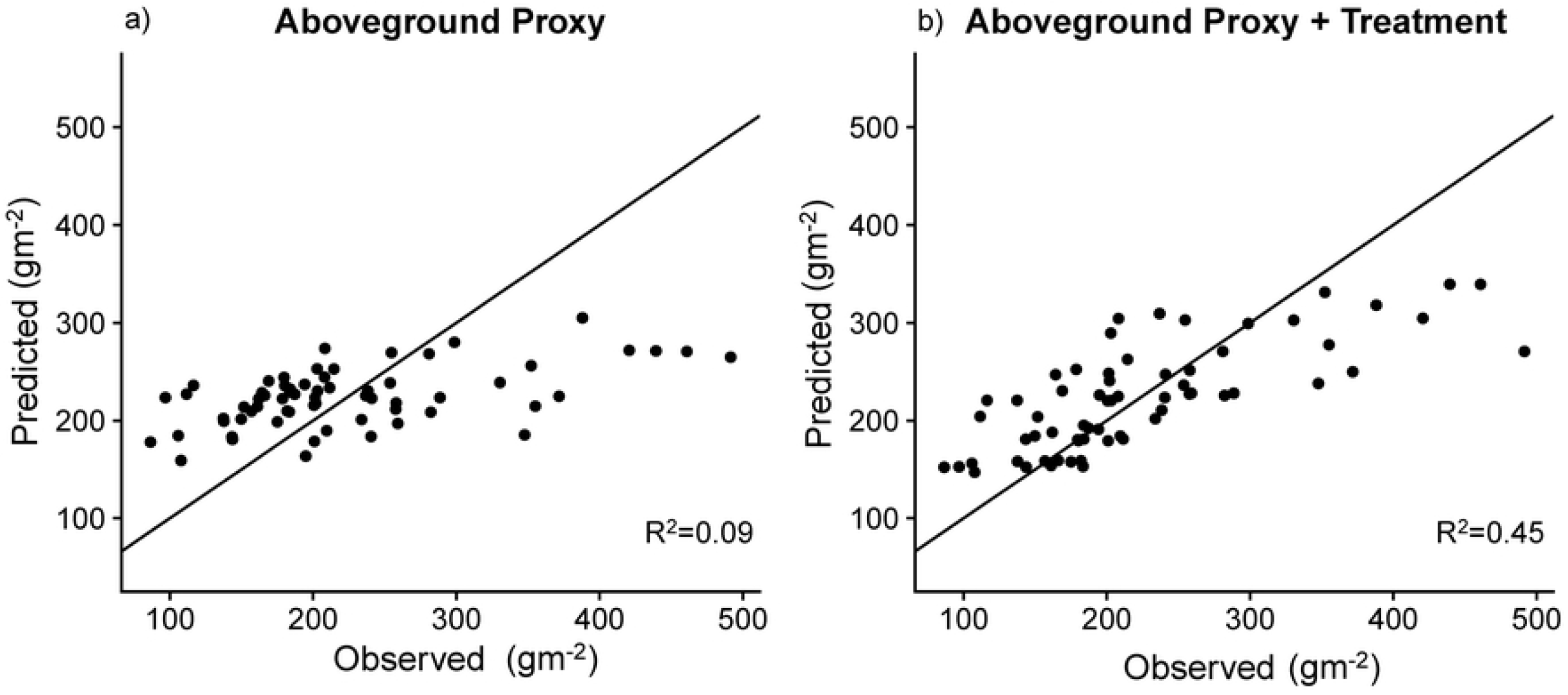
Shoot production does not predict root production. a) Predicted versus observed scatterplot for root production as predicted by shoot production as an aboveground proxy, and b) predicted versus observed scatterplot for root production as predicted by defoliation treatment, cultivar identity, and shoot production. For reference, the 1:1 line of “perfect fit” is plotted along with an in-sample median Bayesian R^2^ for both predictive models.

## Discussion

Severe defoliation resulted in substantially greater shoot production when applied infrequently, but reduced root production among the bahiagrass cultivars. Averaged across all defoliation treatments, root production was also more strongly variable among cultivars than was shoot production. Thus, our results suggest that severe defoliation can trigger a tradeoff between aboveground and belowground allocation in managed subtropical pastures, and that the extent of this tradeoff depends in part on cultivar identity. Contrary to Georgiadis et al. (50) and Briske and Richards (28) who suggested that overcompensation is only likely to occur under water-limitation, or given concomitant fertilization, we found significantly greater shoot production in response to severe defoliation under limited fertility and abundant soil water. Compared with mild defoliation, all cultivars exhibited this compensatory aboveground growth response to severe defoliation, but only when defoliation was applied infrequently (similar to Gates et al. (51)). However, the severe, but infrequent defoliation treatment that led to aboveground compensatory growth also suppressed root production. Thus, under low-input conditions, manipulating defoliation intensity and frequency to enhance forage production could evoke a tradeoff between shoot and root production.. Given the substantial literature demonstrating the importance of root carbon for maintenance of soil carbon pools (17,18,22), these altered allocation patterns may have significant consequences for carbon cycling, and hence soil carbon sequestration services, in managed subtropical pastures. Moreover, use of simple aboveground proxies, such as leaf area/biomass, are unlikely to help constrain predictions of root production over large spatial scales.

Our results differ from the short-term responses measured by Ziter and Macdougall (12) and Hamilton III et al. (52) where a single defoliation event stimulated root production and root exudation, respectively. Moreover, the results reported here appear to conflict with measurements of standing root biomass, root exudation rates, and their connections to microbial biomass and soil carbon, across a system of long-term grazing exclosures on a similar pasture site, as reported in Wilson et al. (22). These discrepancies suggest that root responses to short-term grazing/defoliation events can strongly differ from season-long responses to grazing regimens where both intensity and frequency of defoliation are expected to mediate plant regrowth strategies (28). Moreover, long-term impacts of grazing exclusion in bahiagrass-dominated subtropical pasture appear to involve pronounced phenotypic shifts in root:shoot ratios, whereby absence of grazing favors lower root:shoot ratios, even when holding species composition constant (22)On the other hand, Thornton and Millard (53) found that greater severity of defoliation resulted in lower root mass (but greater N uptake per unit of root mass), which is consistent with our findings. Meanwhile, Dawson et al. (30) report that weekly defoliation over a growing season reduced root biomass compared with no defoliation, but infrequent defoliation (every 8 weeks) had no effect. Our ambivalent findings on the role of frequency of defoliation were thus somewhat surprising. Although we observed marked suppression of variability of production under our severe + frequent treatment (see e.g., Fig 1), root production was not markedly lower than in our severe + infrequent treatment. Overall, it appears that in our system, severity, not frequency, of grazing is the more important determinant of grass root production.

We observed substantial overall variability in root production among the grass cultivars. However, it does not appear possible to predict cultivar-level belowground responses to specific grazing regimens based on observations of aboveground compensatory growth responses. As we hypothesized, the cultivars selected for enhanced upright growth habit (Tifton 9, UF-Riata, (31)) exhibited less overall root production, especially Tifton-9, compared with the widely naturalized decumbent types (Argentine, Pensacola), especially Argentine. On the other hand, all cultivars responded equally negatively to severe defoliation *per se*, and we observed similar total root production among all cultivars in the severe + frequent defoliation treatment, a scenario reasonably representative of overstocked pastures. These results contradict the theory that more grazing-tolerant genotypes, in our case Argentine and Pensacola, will have lower root production as a consequence of greater post-grazing allocation to shoot regrowth (28,30). Instead, it appears that cultivars simply vary in root growth potential, but that severe defoliation, especially when applied frequently, overwhelms this variability.

Contrary to hypothesis, our study revealed that shoot and root production are decoupled at fine spatial scales, at least in our experimental plots, with shoot production explaining only 8% of the in-sample variation in root production. By contrast, defoliation treatment and especially cultivar identity appear to be very important for predicting root production in this system, together accounting for roughly half the observed variance in root production. Gill et al. (14) reported some success in predicting belowground NPP using an algorithm based only on aboveground biomass and climate. However, their model consistently under-predicted root production in more productive sites. Interestingly, we observed a similar severe underprediction of root production in our more productive plots. Thus, we caution against using aboveground proxies to predict belowground production, even within uniform and homogeneous ecosystems, such as the planted pasture system where we worked. Our results suggest that knowledge of grazing management and cultivar identity (in addition to species-level variations in composition, (54,55)) are critical for generating accurate predictions of BNPP. Moreover, half of the variance in belowground production was unexplained, even in our best model, suggesting significant spatial heterogeneity in root system productivity that should be further investigated. Given recent calls highlighting the importance of plant roots to future progress in biogeochemical modeling and the quest to find reliable, scalable aboveground proxies to indirectly infer root processes (36,37), our results are a sobering reminder of the challenges inherent to linking production above and belowground. Accordingly, we suggest that a high priority for future research is to study belowground root-rhizosphere processes using spatially-explicit sampling protocols designed to maximize insight into heterogeneity at various spatial and temporal scales.

At the large scale, McNaughton (1998) (8) found that grazing intensity is uncorrelated with standing root biomass or productivity in the Serengeti. However, in speciose natural grasslands plant diversity may confer a stabilizing influence on root production (55,56). By contrast, monoculture pasture systems may respond more like mesocosm systems where high defoliation intensity is associated with reduced root biomass (24). Moreover, since a large proportion of managed grasslands are dominated by single species, variation in root production among cultivars may represent an especially important component of diversity. Grazing management may need to be matched to cultivar-level characteristics to optimize both forage and root production, and establishment of planted pastures with multiple cultivars or genotypes may be a viable, yet underappreciated, strategy for enhancing functional diversity. For instance, combining upright and decumbent cultivars may introduce beneficial genotypic diversity that could maximize utilization of both above and belowground resources via niche complementarity (57,58). Additionally, cultivar-level variability suggests the potential for ecologists to collaborate with plant breeders to improve the sustainability of grassland agroecosystems by development of improved forage cultivars selected for superior belowground traits.

Overall, our results suggest that intermittent severe defoliation can elicit much greater shoot growth, but have neutral or negative effects on root production. It is possible that a more moderate defoliation intensity than we tested would have led to similar stimulation of aboveground compensation without the negative consequence for root production, a possibility our study was not designed to test. Neither did our study consider impacts of defoliation on rhizome biomass, but we note that our intent was to focus on root production since it appears to be of greater relevance for soil carbon sequestration than other compartments of plant biomass (17). Likewise, it is also possible that the lower fine root production we measured may have been compensated for by greater rhizodeposition/root exudation. However, this possibility seems unlikely given that rates of root exudation generally correlate to fine root surface area (22,59).

## Conclusions

Root production is critical for maintaining and increasing soil carbon pools in grassland ecosystems, yet findings on the immediate and long-term effects of grazing on root production remain variable. We hypothesized that severe defoliation, if applied infrequently, might lead to overyielding of shoots, but would have only small impacts on root production. Moreover, we hypothesized that cultivars selected for an upright growth habit would show less root production overall, and would be more sensitive to defoliation stress. Overall, we found that severe defoliation *per se*, regardless of frequency, suppressed root production, even as infrequently applied severe defoliation increased shoot production. Thus, it appears that manipulating timing and intensity of grazing to optimize forage production might evoke a negative tradeoff with root production. We did find support for the hypothesis that recently developed upright cultivars have lower root production, and a lower root:shoot ratio, than widely naturalized decumbent cultivars. The main limitation of our work is that realistic animal grazing management can differ from experimentally imposed defoliation in two major ways: 1) grazing impacts will fall along a spectrum of timing and intensity with more intermediate values than can be tested in a randomized factorial experiment, and 2) grazers will return a certain fraction of consumed carbon and nutrients in the form of manure and urine, creating heterogeneous patches of varying nutrient availability. Moreover, we also caution that year-year variability in growing conditions can induce variability in experimental effects. Ideally, we recommend long-term (3+ year) field studies of controlled grazing (or defoliation) to begin to properly estimate the random effects of such year-year environmental fluctuations.

In addition to recommending greater future consideration of intraspecific variations in belowground responses to grazing, our work supports the need to perform season-long measures of belowground productivity to obtain reliable estimates of belowground production that can be used to parameterize soil carbon models. Our data also suggest that reliance on aboveground proxies is, unfortunately, not justified at least for subtropical pastures. In addition, given the limitations of observational and comparative work, we suggest that longer-term field manipulations are necessary to evaluate a suite of grazing management scenarios across plant composition treatments. Such experiments will significantly improve our ability to inform the design and management of grassland agroecosystems for meeting aboveground (forage) production goals while also optimizing belowground production, and thus soil carbon sequestration and other soil carbon mediated ecosystem services such as nutrient retention and water cycling (2).

## Acknowledgements

For significant assistance with field work and data collection we thank Carly Althoff, Jessica Wilson, Trevor Caughlin, Anand Roopsind, James Estrada and Bryan Tarbox.

## SUPPORTING INFORMATION

**S1 Fig. Diagram showing layout of plots**. North is top of the page. Legend: Defoliation severity-Red = Severe Defoliation (5 cm), Blue = Lenient Defoliation (15 cm). Defoliation frequency - 2wk = Defoliated every 2 weeks, 4wk = Defoliated every 4 weeks. Bahia cultivar identity - A = Argentine, P = Pensacola, T9 = Tifton 9, R = UF-Riata.

**S1 Table. Meteorogical data from our study site during study season**. Ona Range Cattle Research and Education Center. Accessed from the Florida Automated Weather Network (FAWN), http://fawn.ifas.ufl.edu/.

